# LMethyR-SVM: Predict human enhancers using low methylated regions based on weighted support vector machines

**DOI:** 10.1101/054221

**Authors:** Jingting Xu, Hong Hu, Yang Dai

## Abstract

**Background:** The identification of enhancer is a challenging task. Various types of epigenetic information including histone modification have been utilized in the construction of enhancer prediction models based on a diverse panel of machine learning models. However, DNA methylation profiles generated from the whole genome bisulfate sequencing (WGBS) have not been fully explored for their potential in enhancer prediction despite the fact that low methylated regions (LMRs) have been implied to be distal active regulatory regions.

**Method:** In this work we propose a prediction framework, LMethyR-SVM, using LMRs identified from cell-type-specific WGBS DNA methylation profiles based on an unlabeled-negative learning framework. In LMethyR-SVM, the set of cell-type-specific LMRs is further divided into three sets: reliable positive, like positive, and likely negative, according to their resemblance to a small set of experimentally validated enhancers in the VISTA database based on an estimated non-parametric density distribution. Then, the prediction model is trained by solving a weighted support vector machine.

**Results:** We demonstrate the performance of LMethyR-SVM by using the WGBS DNA methylation profiles derived from the H1 human embryonic stem cell type (H1) and the fetal lung fibroblast cell type (IMR90). The predicted enhancers are highly conserved with a reasonable validation rate based on a set of commonly used positive markers including transcription factors, p300 binding and DNase-I hypersensitive sites. In addition, we show evidence that the large fraction of LMethyR-SVM predicted enhancers are not predicted by ChromHMM in H1 cell type and they are more enriched for the FANTOM5 enhancers.

**Conclusion:** Our work suggests that low methylated regions detected from the WGBS data are useful as complementary resources to histone modification marks in developing models for the prediction of cell type-specific enhancers.

## Introduction

Enhancers play an important role in temporal and cell type-specific activation of gene expression [1]. Enhancers are short in their length and often consist of clusters of binding sites for transcription factors (TFs) [2]. The identification of enhancers is challenging both experimentally and computationally due to their distal locations from the transcription starting sites (TSSs) of the target genes [3]. There have been reported large-scale experimental approaches to enhancer discovery for human heart tissue [4]. The VISTA database reported 2,316 *in vivo* validated enhancers for mouse and human [5] (as of 06/09/2015). In addition, the FANTOM 5 CAGE gene expression atlas identified and characterized 43,011 enhancer candidates across the majority of human cell types and tissues from the 432 primary cell, 135 tissue and 241 cell type samples [6]. However, the number of validated enhancers (32,693 as of 04/28/2016) is far from the estimated 500,000-1 million enhancers in the human genome [7], suggesting the importance of enhancer prediction using of computational models.

The activity of enhancers has been shown to correlate with certain chromatin properties; such as accessibility of DNA to which TFs are bound and the post-translational modification in the tails of histone proteins in chromatin in the vicinity of the active enhancers [8–12]. Previous work has explored this information to build models for the prediction of enhancers [13–20]. Commonly, p300 sites overlapping with DNase-I hypersensitivity sites (DHSs), the sites marked by H3K4me1, H3K4me2, H3K4me3 and H3K27ac, are the most explored information in the model development [13, 14, 16, 17, 21, 22]. p300, a transcriptional co-activator recruited by TF complexes, has been found at a large number of enhancers. DHSs are the accessible chromatin regions that are functionally related to transcriptional activity. While these methods successfully identify many enhancers, the predicted enhancers tend to be bound by the known p300 binding sites. Thus, they may potentially miss out other types of enhancers. On the other hand, the analysis of whole genome bisulfate sequencing (WGBS) DNA methylation profiles has led to the identification of low methylated regions (LMRs) [23, 24]. LMRs are shown to be associated with distal regulatory elements, as they are enriched for active histone marks (e.g. H3K4me1), DHSs, p300 and TF binding sites (TFBSs) [23–26]. However, the potential of DNA methylation data for enhancer prediction has not been fully explored.

Herein, we systematically assess the usefulness of LMRs for enhancer prediction. Three questions are specifically addressed: (i) Can LMRs be used as putative enhancer sequences to build a prediction model? (ii) How is the model compared to those using information derived from p300 binding sites, DHSs, and histone modification marks? (iii) Are the predicted enhancers cell-type specific? To this end, we propose a novel framework, called LMethyR-SVM, which builds a model for enhancer prediction from sequences of cell-type-specific LMRs based on a weighted support vector machine. Although LMRs are enriched for marks of active regulation, not all sequences of LMRs are necessarily enhancers. For this reason, LMRs are initially called unlabeled. To dissect the LMRs, *in vivo* validated enhancer sequences from the VISTA enhancer database [5] are explored to divide the unlabeled set into a reliable positive set, a likely positive set and a likely negative set, representing our confidence in the LMRs being putative enhancers. By encoding sequences based on a *k*-mer scheme, this division is achieved according to the density ranks of each LMRs on a density distribution estimated from the LMRs that overlap with the VISTA enhancers. With a random negative set, the weighted support vector machine (wSVM) is then designed to learn a prediction model by best dividing the unlabeled set of the LMRs into three datasets, namely, reliable positive, likely positive and likely negative. LMethR-SVM trains on sequences of cell-type-specific LMRs and predicts enhancers by scanning the genome without using information of DNA methylation. We compare the performance of LMethyR-SVM with existing methods using the WGBS data from the H1 human embryonic stem cell type (H1) and the fetal lung fibroblast cell type (IMR90). We further investigate the cell-specificity of the predicted enhancers.

## Materials and Methods

Our framework, LMethyR-SVM, begins with a DNA methylation profile obtained from a specific cell type. Then the cell-type-specific LMRs are identified; the set of unlabeled LMRs and the VISTA enhances are used to build an estimated density distribution in the *k*-mer space. Then, each LMR sequence is assigned a rank based on its density. Finally, the wSVM learns a model from the unlabeled LMRs and a random dataset by determining the best division of the unlabeled LMR set into a reliable positive (RP) set, a likely positive (LP) set, and likely negative (LN) set. The proposed framework of LMethyR-SVM is shown in **Fig 1**.

**Fig 1.**
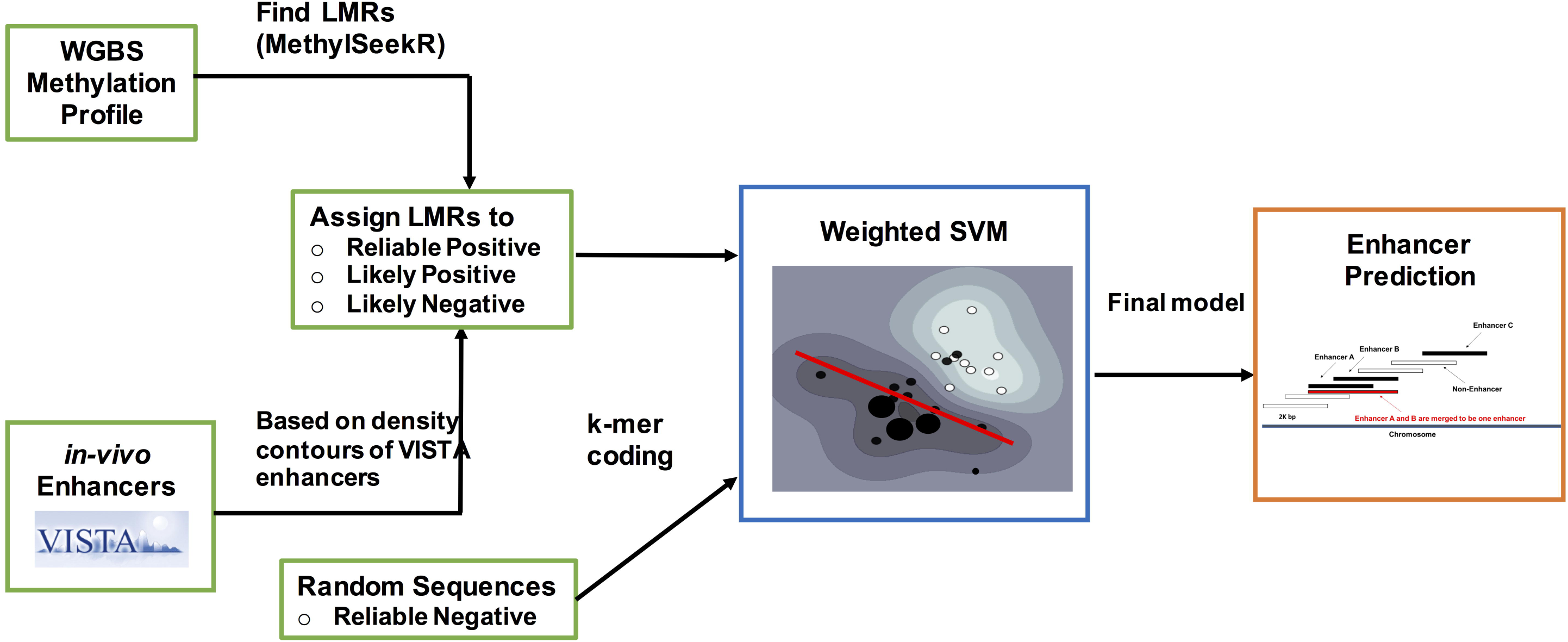
The proposed framework of LMethyR-SVM.

### Low methylated regions (LMRs) in H1 and IMR90 cell types

The WGBS data of DNA methylation in H1 and IMR90 cell types were available from the website of the Salk Institute for Biological Studies [27]. The WGBS data were generated using the MethylC-seq technique and the raw data have been mapped and quantified for the methylation level at individual cytosines in the genome. The downloaded data are processed methycytosine profiles. We lifted the data to the UCSC hg19 assembly using liftOver provided at the USCS Genome Browser [28]. LMRs were computed using the Bioconductor package MethylSeekR [24] with the recommended parameters. MethylSeekR identifies hypomethylated regions as stretches of consecutive CpGs with methylation levels below a fixed threshold and further divides them into unmethylated and low methylated regions. Genome regions with SNPs, and X and Y chromosomes were excluded from the analysis since the LMRs could not be reliably detected. More details about this procedure can be found in the previous publications [24, 25].

### The VISTA enhancers

The VISTA Enhancer Browser is a central resource for experimentally validated human and mouse noncoding fragments with gene enhancer activity in transgenic mice [5]. The 967 validated human enhancers were downloaded from the VISTA database (as of 6/09/2015). The VISTA enhancers are highly conserved in other vertebrates or rich for epigenomic evidence (ChIP-Seq) of putative enhancer marks. Although these enhancers are not specific to a cell type, the subsets that overlap with the LMRs from H1 cell type and IMR90 were identified and used as putative cell-type-specific enhancers to estimate an underlying density distribution of putative enhancers. The density distribution for each cell type was used to assign a density rank to each sequence in a set of LMRs.

### Weighted support vector machine (wSVM) for model learning from unlabeled and negative sets

As suggested in the previous studies, LMRs are indicative for distal active regulatory regions that may include enhancers [23–26]. However, the functional role of LMRs in gene regulation is unclear. Since they may also include some sequences that are not relevant to enhancers, the set of LMRs is called unlabeled (U), instead of being labeled as a positive set of enhancers. From this unlabeled set, the LMRs were further divided into three sets: RP, LP, and LN according to their ranks of density, which indicate the degree of resemblance to the LMRs that overlap with the VISTA enhancers for each cell type (the procedure described in next section). To facilitate machine learning, a reliable negative (RN) set of random of sequences was constructed by randomly selecting loci that do not include the sequences from the UCSC exons and the LMRs. The shuffle function in BEDTools was used to randomly permute the genomic locations of a feature BED file (i.e., LMRs) on the human genome (hg19). The tool excludes loci of exons and LMRs while generating a set of sequences that preserve the distribution of length and number on each individual chromosomes as that of the LMRs. In addition, the random sequences containing base ‘N’ besides ‘ATGC’ were excluded. All sequences were represented using a *k*-mer encoding scheme, i.e., each sequence is encoded by a vector *x* with 4^*k*^ entries, each of which indicates the number of times that a particular *k*-mer appears in the sequence. In this work, we used *k*=5 based on our preliminary investigation in wSVM training. Furthermore, each vector *x* was normalized by dividing the sum of all its entries in *x*.

To reflect our confidence in the given sets of RP, LP, and LN of LMRs, the wSVM model was proposed as the following optimization problem.

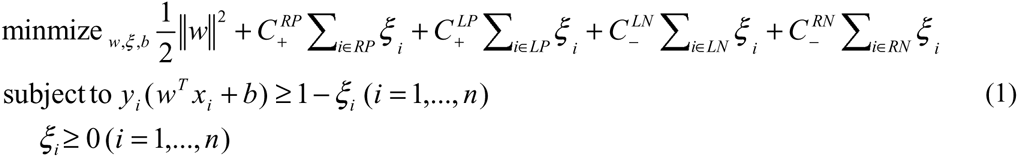
where *ξ*_*i*_ is a slack variable that allows for error for misclassification, and *n* is the total number of training examples. For each *x*_*i*_, it is labeled as *y*_*i*_ = 1 if it is in RP or LP; and *y*_*i*_ =‒1 otherwise. The weights 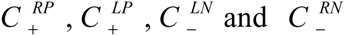 are weight parameters for wSVM to penalize misclassified training examples in RP, LP, LN, and RN, respectively. However, the division of LMRs along with the weights have to be learned based on a cross-validation procedure described below. We restrict 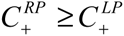 since we have more confidence in the reliable positive set RP than in the likely positive set LP. Consequently, a sequence from RP receives a larger penalty than a sequence from LP if it is classified as negative. Similarly, we set 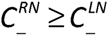 for the reason that we have more confidence in RN than in LN and misclassification of a sequence from RN as positive receives a larger penalty than if a sequence from LN is misclassified as positive class.

### Partition of an unlabeled set of LMRs based on ranks of the LMRs on a density distribution

The following scheme was used to determine our confidence for each sequence in the unlabeled set of the LMRs. The LMRs overlapping the VISTA enhancers were considered as putative enhancers in that cell type. This subset of LMRs was used to generate an estimated underlying density distribution of the sequences in the *k-mer* space. The overlapping VISTA enhancer sequences themselves were not used because they contain primers of several hundred base pairs on flank regions, which may bring “noise” into model training. The R package pdfCluster [29] was used to detect the underlying cluster structure of the sequences based on the kernel density estimation. In contrast to traditional clustering methods based on metric of distance or dissimilarity, the density-based clustering method associates cluster(s) with the regions around the modes of the density underlying the data [30]. By measuring density of a LMR to its assigned density center in the *k-mer* space, the sequence was given a rank. The higher its rank is, the closer a sequence is to its cluster center. LMRs that were ranked at the top *δ* quantile were labeled as RP; those under 15% quantile were labeled as LN; and the remainder were labeled as LP. The value of *δ* was learned through the cross-validation (CV) procedure in wSVM training (more detail in Supporting Information **S1 File**).

### Protocol for training and testing

The wSVM model described above can be solved as an optimization problem if a quantile threshold *δ* for dividing the LMRs into RP and LP, and a set of weights 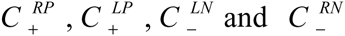 are given. To determine the best values for these parameters, a 5-fold CV procedure was used to choose a group of models with the highest cross-validated *F-*scores. The parameter set that generated the model with the highest precision from the group was chosen as the best parameter set. Here *F*-score is defined as

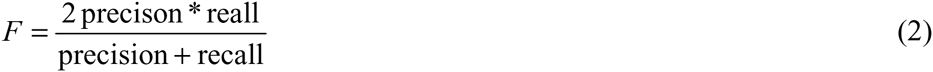

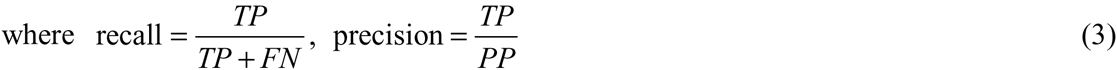
and *TP* and *FN* are the numbers of LMRs in RP and LP that are predicted as positive and negative, respectively; and *PP* is the number of LMRs that are predicted as positive. Note that the definition of *F*-score seems to be identical to the ones used in standard binary learning. However, it is different in the sense that is dependent on the RP and LP sets, which are determined by the value of quantile threshold *δ*. Therefore, the best model cannot be chosen merely based on *F*-score, rather determined by the one with highest precision among the models with the best F-scores. In addition, genomic coverage, as an additional constraint in model selection was required. Genomic coverage is defined as the fraction of nucleotides that are covered by the predicted enhancers in the hg19 human genome. In the training process, the genomic coverage of the model’s prediction on Chromosome 1 was used as a proxy for the entire genome. In the CV process, models that predict enhancers on Chromosome 1 with genomic coverage more than a prescribed threshold value were excluded. With this additional constraint, we could effectively avoid generating models that predict a large fraction of the genome as enhancers. The details of parameter tuning and the cross-validation procedure can be found in the Supporting Document **File S1**, **S1 Fig** and **S2 Fig**). Libsvm [31] was used to train wSVMs with linear kernels during the CV process and for the final prediction model for each cell type, and the heuristic procedure was implemented using shell scripts.

The enhancer prediction was carried out by sliding a window of size 2,000 bp for every 500 bp on the human genome according to the score assigned to each window by the final wSVM model. Windows with positive scores were predicted as enhancer windows. Overlapping enhancer windows were joined and the covered region was reported as one enhancer; the covered region may be reported for more than one enhancer by separating at any possible negatively scored windows in it, a situation can occur by the way of window sliding.

### Validation procedure for predicted enhancers

The publically available data on epigenetic marks associated with enhancer activities were used in validation according to the set of the same criteria as employed in the previous work [13]. A true positive marker (TPM) is defined as any of the DHS, CBP/p300 sites and enhancer-associated TFs. The enhancer windows are classified as “validated”, “misclassified” or “unknown” as follow.

1. If the nearest TPM lies within 1,000 bp of a predicted enhancer windows and the nearest TSS (we use the UCSC TSSs) is greater than 1,000 bp away from the TPM, then the enhancer is “validated”.
2. If a TSS lies within 2,500 bp of the predicted enhancer window, and the nearest TPM is either greater than 1,000 bp away from the enhancer or within 1,000 bp of the TSS, the enhancer is “misclassified”.
3. If there is neither TPM within 1,000 bp nor TSS within 2,500 bp of the enhancer, it is “unknown”.

The “validated” enhancers were divided into 6 mutually exclusive states as in [13]: “p300+/-DHS”, “DHS only”, “TF+DHS”, “TF only”, “TF+p300”, “p300+DHS+TF”. For example, “p300+DHS+TF” means that an enhancer window is validated by all the three markers; “p300+/-DHS” includes “p300+DHS” and “p300”. The percentages of the predicted enhancer windows that were classified as “validated”, “misclassified” and “unknown” were calculated.

The experimental data of DHSs and p300 binding sites for the two cell types, and enhancer-associated TFs, i.e., NANOG, CEBPB, TEAD4 for H1 and CEBPB for IMR90, were downloaded from “ChIP-Seq Experiment Matrix” from the ENCODE website [32]. The p300 binding sites for IMR90 were obtained from the previous publication RFECS [13]. The “uniform”, or “narrow peak” type of the ChIP-Seq data for each marker was selected. These types of data were consolidated from multiple experiment results and were controlled at a significantly small false positive rate. Meanwhile, since the total length/coverage of the sites are small, the validation by overlapping these markers is unlikely due to randomness.

### The FANTOM 5 enhancers

The 32,693 enhancers reported from FANTOM5 – the Functional Annotation of the Mammalian Genome project were downloaded from the consortium website (as of 04/28/2016) [33]. The *in vivo* active enhancers are identified based on the distinct bidirectional CAGE (Cap Analysis of GENE Expression) pattern using the FANTOM5 panel of tissue and primary cell samples [6]. FANTOM5 CAGE expression atlas includes 43,011 enhancer candidates in 432 primary cell, 135 tissue, and 241 cell type samples from human. Although both H1 and IMR90 cell types are not included, the FANTOM enhancers are used as a proxy of active enhancers to check the extent of enrichment in the predicted enhancers. A predicted enhancer window is considered to be overlapping with a FANTOM enhancer if the overlap is at least by 1 bp.

### Statistical tests

All statistical test results for proportional difference and enrichment were obtained using z-test in R.

## Results

As described previously, MethylSeekR was used to identify LMRs from H1 and IMR90 WGBS methylation profiles. It found 29,138 and 37,033 LMR sequences for H1 and IMR90, respectively. The median length of LMRs is 410 bp in H1 and 622 bp in IMR90. As expected, they mostly reside in distal regions that are more than 2,000 bp away from TSSs (**Fig 2 (a)**). In addition, only 9.2% and 44.2% of LMRs overlapped with the p300 binding sites in H1 and IMR90, respectively. The higher percentage of the overlap in IMR90 may be due to a much larger number of p300 binding sites (52,878) in comparison to that in H1 (8,934). The LMR sequences with length less than 200 bp or greater than 3,000 bp were removed for subsequent model training, resulting in 23,017 and 34,174 LMR sequences in H1 and IRM90, respectively.

**Fig 2.**
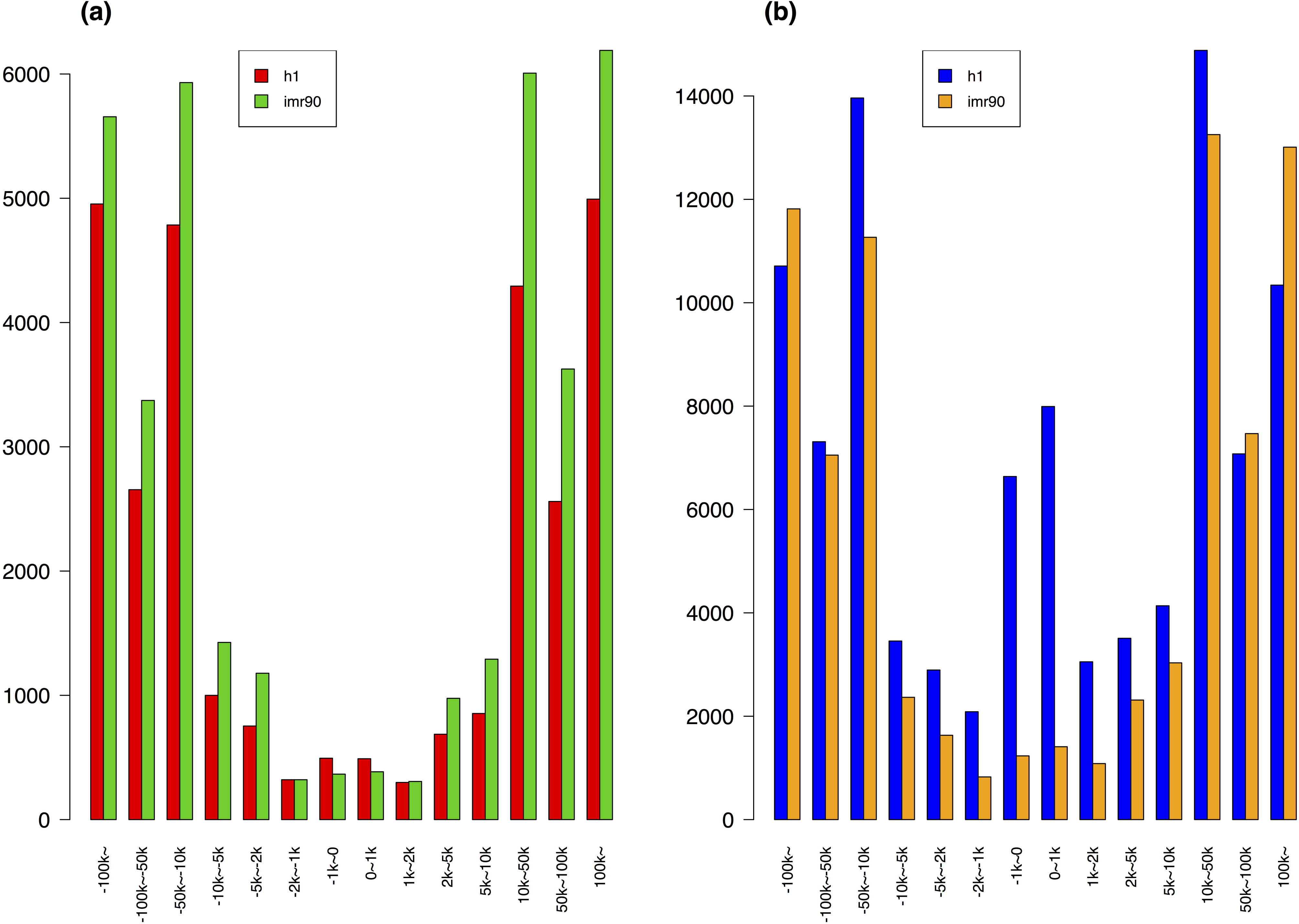
The distributions of the LMRs and predicted enhancer windows. (a): The distributions of the LMRs to the nearest TSSs; red for H1 and green for IMR90. (b): The distribution of all the enhancer windows to their nearest TSSs; blue for H1 and brown for IMR90. The distance was measured from the center point of a sequence to its nearest TSS.

We found that 106 LMRs overlap with 105 VISTA enhancers for H1 and 129 LMRs overlap 117 Vista enhancers for IMR90 by at least 1 bp. These LMRs are called the VISTA-overlapping LMRs. Next, we applied pdfCluter to derive an estimated nonparametric density distribution of the VISTA-overlapping LMRs in each cell type and then assigned each LMR a rank according to its density. Then, the 5-fold CV procedure was applied to find the best quantile cutoff to divide the LMRs into sets of RP, LP, and LN, and the corresponding weights in wSVM for each cell type (Supporting document **File S1**). We chose genomic coverage at 5% for model selection, although models 10% and 15% of genomic coverage were also developed. The value of 5% was determined because the number of the predicted enhancers with this constraint generated a similar number of enhancers as other methods so that a fair comparison could be made. The quantile cutoffs between RP and LP learned from wSVM were both 95% for H1 and IMR90.

A summary of the predicted enhancers is given in **Table 1**. The distributions of the genomic locations of all the enhancer windows with respect to the nearest TSSs are shown in **Fig 2 (b)**). It can be observed that the majority of them are distal to the nearest TSSs. Interesting, the distribution remained almost the same even after removing the LMR sequences overlap with promoter regions defined as 2000 bp up- and down- stream of the TSSs (S3. Fig in Supporting document **File S1**).

**Table 1.** The summary of the predicted enhancers obtained from LMethyR-SVM. ^(a)^ Enhancers are the continuous regions obtained by concatenating all overlapping enhancer windows and removing regions covered by non-enhancer windows. ^(b)^ The genome coverage is the estimate from Chromosome 1.

|  | Number of enhancer windows | Number of enhancers<sup>(a)</sup> | Estimated Genomic Coverage<sup>(b)</sup> | Median length of enhancers | Maximum Length of enhancers |
| --- | --- | --- | --- | --- | --- |
| H1 | 98,045 | 34,437 | 3.67% | 2,500 bp | 66,000 bp |
| IMR90 | 77,762 | 35,203 | 3.05% | 2,500 bp | 113,000 bp |

<sup>(b)</sup> The genome coverage is the estimate from Chromosome 1.

The numbers of the enhancer windows are 98,045 and 77,762 in H1 and IMR90, respectively. We obtained the predicted enhancers by concatenating overlapping enhancer windows. This resulted in 34,437 and 35,203 enhancers in H1 and IMR90, respectively. Since some of the predicted enhancers are quite long, e.g. the longest enhancer in IMR90 is of 113,000 bp long, we further selected the highest-scored enhancer window from each enhancer as the representative of this enhancer for fair comparison with other methods.

### The performance of LMethyR-SVM

Since there is no true set of cell-type-specific enhancers for validation, we evaluated the predicted enhancers from three aspects. We first compared our results with those obtained from three state-of-art enhancer prediction methods: ChromHMM [20], RFECS [13], and EnhancerFinder [16]. We also used a random set of sequences to serve as a baseline for comparison, which was generated by randomly selecting 2,000 bp windows to be enhancers with the same number as ours. The three methods utilize various types of chromatin modification information coupled with supervised or unsupervised machine learning models. However, none of them has used epigenetic modification information of DNA methylation. Since it would be ideal for fair comparison if the lengths of predicted enhancers were similar, we chose the highest-scored enhancer windows in this validation analysis. The results of the predicted enhancers at genomics coverage 5% based on the validation procedure using TPMs are shown in **Fig 3**. Note that for IMR90, RFECS is the only tool that has made prediction besides LMethyR-SVM. In general, a high percentage of “validated” and low percentages of “misclassified” and “unknown” suggest high confidence in a prediction model. These criteria allow the performance comparison among different prediction models with relatively small bias if a common set of experimental markers is used.

**Fig 3.**
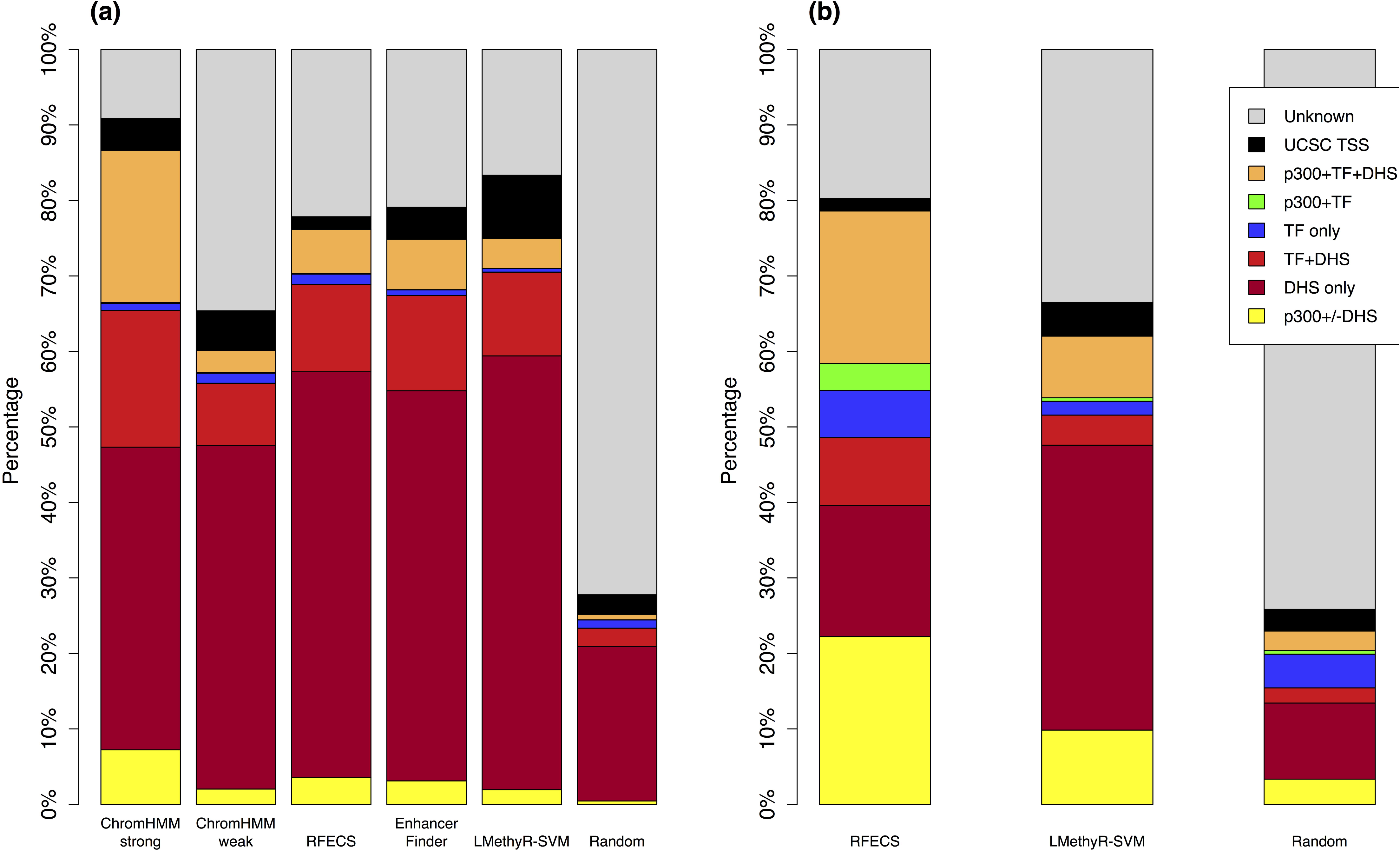
Results of comparison with other enhancer prediction models. (a) for H1 and (b) for IMR90. “Validation” rates were measured as percentages of overlaps with either DHSs, p300 sites or cell-type-specific transcription factor binding sites (NANOG, CEBPB and TEAD4 for H1 and CEBPB for IMR90); “Misclassification” rates were measured as percentages of overlaps with the UCSC annotated TSSs. “Validated” enhancers can be further divided into one of the mutually exclusive categories: “p300+/-DHS”, “DHS only”, “TF+DHS”, “TF only”, “TF+P300”, “p300+DHS+TF”. For LMethyR-SVM, the highest-scored enhancer windows were used. The total numbers of the enhancers in H1 predicted from the individual methods are 17,828 (ChromHMM Strong), 217,350 (ChromHMM weak), 54,121 (RFECS), 37,263 (EnhancerFinder), 34,437 (LMethyR-SVM) and 34,437 (Random). The total numbers of the enhancers in IMR90 predicted from the individual methods are 82,392 (RFECS), 35,203 (LMethyR-SVM) and 35,203 (Random).

ChromHMM [20] used a hidden Markov model to segment the genome of 200 bp windows into multiple states on the basis of consensus on patterns. Then, domain experts manually annotated each state. ChromHMM integrated most of the existing ChIP-Seq data on chromatin modifications. It predicted enhancers for each of the 9 different cell types: H1, GM12878, Hepg2, Hmec, Hsmm, Huvec, K562, Nhek, and Nhlf. These data are available from the Annotation Database at the UCSC Genome Browser website. For H1 cell type, ChromHMM predicted 235,178 enhancers, which are further catalogued into either 217,350 “weak enhancers” or 17,828 “strong enhancers”. Since it used the ChIP-Seq data from ENCODE for model learning, it is not surprising that ChromHMM is one of the best performers, especially for its “strong enhancer” category (**Fig 3 (a)**). Its validation rate (86.65%) is significantly higher than that of LMethyR-SVM (74.87%, *P* < e-15). However, its “weak enhancers” category has far too many enhancers, resulting in a worse validation rate (60.14%, *P* < e-15) compared to LMethR-SVM (74.87%).

The second method, RFECS, was trained with a random forest learning model on 24 ChIP-Seq histone modification data from ENCODE and the p300 binding data for IMR90 generated from their own study. The best model predicted enhancers with the use of the three key histone modifications. RFECS makes prediction by sliding windows of 1,500 bp - 2,500 bp. It has predicted 54,121 enhancers for H1 and 82,392 for IMR 90 [13]. However, the published enhancers are in the form of a 1bp locus. Therefore, we expanded this locus by 1,000 bp on both sides for fair comparison. The validation rate of the RFECS prediction is 76.14% for H1, indicating that LMethyR-SVM has a lightly worse performance (74.94%, *P* =2.6e-5) (**Fig 3 (a)**). Similarly, RFECS for IMR90 has a validation rate of 78.60%, which is better than LMethyR-SVM (62.02%, *P* < e-15) (**Fig 3 (b)**). The performance of RFECS could be due to its extensive training based on the comprehensive ChIP-seq data.

The third method, “EnhancerFinder”, uses a multiple kernel learning approach to integrate DNA sequence motifs, evolutionary patterns, and 2,469 functional genomics datasets generated by the ENCODE project and smaller scale studies [16]. The tool predicted enhancers for multiple cell types (including H1) as well as multiple organs, but not for IMR90. The published 82,490 enhancers have a validation rate of 74.87%, which is not significantly different compared with LMethyR-SVM (74.94%, *P* =0.41) (**Fig 3 (a)**).

In summary, the performance of LMethR-SVM evaluated using TPMs is highly encouraging, considering the fact that only DNA methylation data were used in building the prediction model and all the other methods relied on the abundant ChIP-Seq experiment data of histone modifications, and sites of p300 binding and DHSs. The way of training and evaluation by all the other models could favor towards the validation of their results and may lead to overestimated performance. On the other hand, the experiment data of the DNA methylation used in LMethyR-SVM is independent of any of the ENCODE experiment results. This suggests that LMRs is informative as a source to build enhancer prediction models.

Next, we set out to examine the extent of overlaps between the predicted enhancers made by LMethyR-SVM and ChromHMM for H1; LMethyR-SVM and RFECS for IMR90. We used ChromHMM for H1 since it can be considered as “truth” on the basis that its prediction was made on the analysis of the most comprehensive ENCONE datasets. In addition, we also analyzed the overlap with the FANTOM5 enhancers. The results are summarized in **Table 2**. This part of the analysis was performed only on the predicted enhancers that were validated with the TPMs. From Table 2 we observe that 11.79% of the enhancer windows predicted by LMethyR-SVM in H1 overlap with the FANTOM5 enhancers, which is significantly higher than that of the ChromHMM enhancers (4.67%, *P* < e-15). Similarly, 14.24% of the enhancer windows predicted by LMethyR-SVM in IMR90 overlap with the FANTOM5 enhancers, representing a significantly higher proportion than that of RFECS (13.78%, *P* =0.013). This result for IMR90 is interesting because RFECS has a better performance than LMethyR-SVM in terms of the validation rate based on the TPMs in IMR90. Further, we took the enhancer windows that are uniquely predicted by each method and compared the enrichment for the FANTOM enhancers. It can be seen that 10.01 % of the enhancer windows uniquely predicted from LMethyR-SVM in H1 overlaps with the FANTOM5 enhancers, representing a significantly higher proportion than that of ChromHMM (4.12%, *P* < e-15). However, the enhancer windows uniquely predicted from LMethyR-SVM in IMR90 have a lower proportion (8.63%) for the FANTOM5 enhancers compared to that of RFECS (12.35%, *P* < e-15). Finally, the overlapped enhancer windows generated from the two corresponding methods in each cell type always included a significantly higher number of the FANTOM5 enhancers compared to the enhancers unique to each individual method (data not shown). This analysis provides further evidence indicating that LMRs may provide a complementary sequence profile other than the regions marked by histone modification, p300 binding and DNSs sites for enhancer prediction.

**Table 2.** The summary of the overlap between the predicted enhancers and the FANTOM5 enhancers. ^(a)^ Unique: the number of the enhancer windows that were uniquely predicted by a method and validated by TPMs. ^(b)^ Total: the total number of the enhancer windows that were predicted by a method and validated by TPMs. ^(c)^ %FANTOM: the percentage of the enhancer windows that overlap with the FANTOM5 enhancers.

|  |  | Unique <sup>(a)</sup><br>(%FANTOM<sup>(c)</sup>) | Total <sup>(b)</sup><br>(%FANTOM) |
| --- | --- | --- | --- |
| H1 | ChromHMM | 120,539<br>(4.12%) | 146,172<br>(4.67%) |
|  | LMethyR-SVM | 45,629<br>(10.01%) | 75,947<br>(11.79%) |
| IMR90 | RFECS | 55,665<br>(12.35%) | 64,765<br>(13.78%) |
|  | LMethyR-SVM | 32,775<br>(8.63%) | 51,146<br>(14.24%) |

Finally, we investigated if the LMethyR-SVM predicted cell-type-specific enhancers. This task is challenging because no complete set of validated enhancers exist for each cell type. Therefore, we evaluated the cell-type-specificity indirectly by comparing the LMethyR-SVM predicted enhancer windows for H1 with those predicted by ChromHMM for H1 and 8 other cell types (H1, GM12878, Hepg2, Hmec, Hsmm, Huvec, K562, Nhek and Nhlf). Our argument is that if the overlap between the LMethyR-SVM and ChromHMM predicted enhancers in H1 is higher than those between the LMethyR-SVM for H1 and ChromHMM for other cell types, it may suggest more cell-type-specific enhancers were predicted by LMethyR-SVM. Indeed, it is observed that the LMethyR-SVM predicted enhancer windows have the highest percentage of overlaps (39.2%) with the ChromHMM predicted enhancers windows for H1 cell type (**Table 3**). The proportion of overlap in H1 cell type is significantly higher than that in any of the other 8 cell types (26.5% - 37.6%, P<e-15).

**Table 3.** The comparison of the number of enhancer windows in each cell type predicted by ChomeHMM and the number of windows that overlapped with the enhancer windows predicted by LMethyR-SVM in H1 cell type. ^(a)^ The number of enhancer windows predicted by LMethyR-SVM in H1 overlapped with those predicted by ChromHMM.

| Cell type | # enhancers predicted by ChromHMM | # overlaps between LMethyR-SVM and ChromHMM enhancers <sup>(a)</sup> | Percent of overlap |
| --- | --- | --- | --- |
| H1 | 235,178 | 38,397 | 0.392 |
| GM12878 | 236,344 | 25,934 | 0.265 |
| Hepg2 | 201,190 | 29,365 | 0.300 |
| Hmec | 283,718 | 36,805 | 0.376 |
| Hsmm | 267,423 | 30,743 | 0.314 |
| Huvec | 228,423 | 30,347 | 0.310 |
| K562 | 242,306 | 32,598 | 0.332 |
| Nhek | 272,728 | 28,908 | 0.295 |
| Nhlf | 235,293 | 27,493 | 0.280 |

### The evolutionary conservation in the predicted enhancers by LMethyR-SVM

Enhancers that are active during the early mammalian developmental stage tend to be conserved [34]. H1 cell type is one of the embryonic stem cells (ES cells) derived from the inner cell mass of a blastocyst, an early-stage pre-implantation embryo. In addition, most of the VISTA enhancers are ultra-conserved non-coding DNA sequences on the vertebrate genome when first being selected for testing. Indeed, 99.8% of the VISTA enhancers overlap at least 1 phastCons46ways conserved regions at vertebrate level. PhastCons is a computational tool that finds and annotates conserved segments on the human genome using comparative genomic information. The conserved information was downloaded from the track on the UCSC website [28]. We wanted to check if our prediction model, which uses information on the VISTA enhancers, reflects this inclination of conservation. Since there are about 5 million conserved segments annotated by PhastCons on the entire human genome, they are densely populated, a window of 2,000 bp in length can easily include one conserved element by chance. To avoid over-estimation, each predicted enhancer window was represented by its midpoint and any enhancer window overlapping with exons has been removed for this analysis. If the midpoint locates on any conserved segment, then the enhancer window is considered “conserved”. The random set was also included to serve as the baseline for comparison.

It is shown that for H1 cell type the predicted enhancers obtained from LMethyR-SVM have the second highest proportion of enhancers that overlap at least one conserved segment (**Fig 4**(a)), only after EnhanerFinder, which also has integrated the VISTA enhancer information. ChromHMM predicted the most heterogeneous type of enhancers because its non-supervised model was built based on data from multiple cell types. It shows moderate but above the conservation level of random. There is no significant difference in conservation between its “strong enhancers” and “weak enhancers” (data not shown). For IMR90 cell type, the RFECS predicted enhancers are slightly conserved than that of LMethyR-SVM (**Fig 4 (b)**). Together, LMethyR-SVM predicts the enhancers that tend to be more conserved in H1 cell type, which may be more relevant to the early developmental stage in H1 cell type.

**Fig 4.**
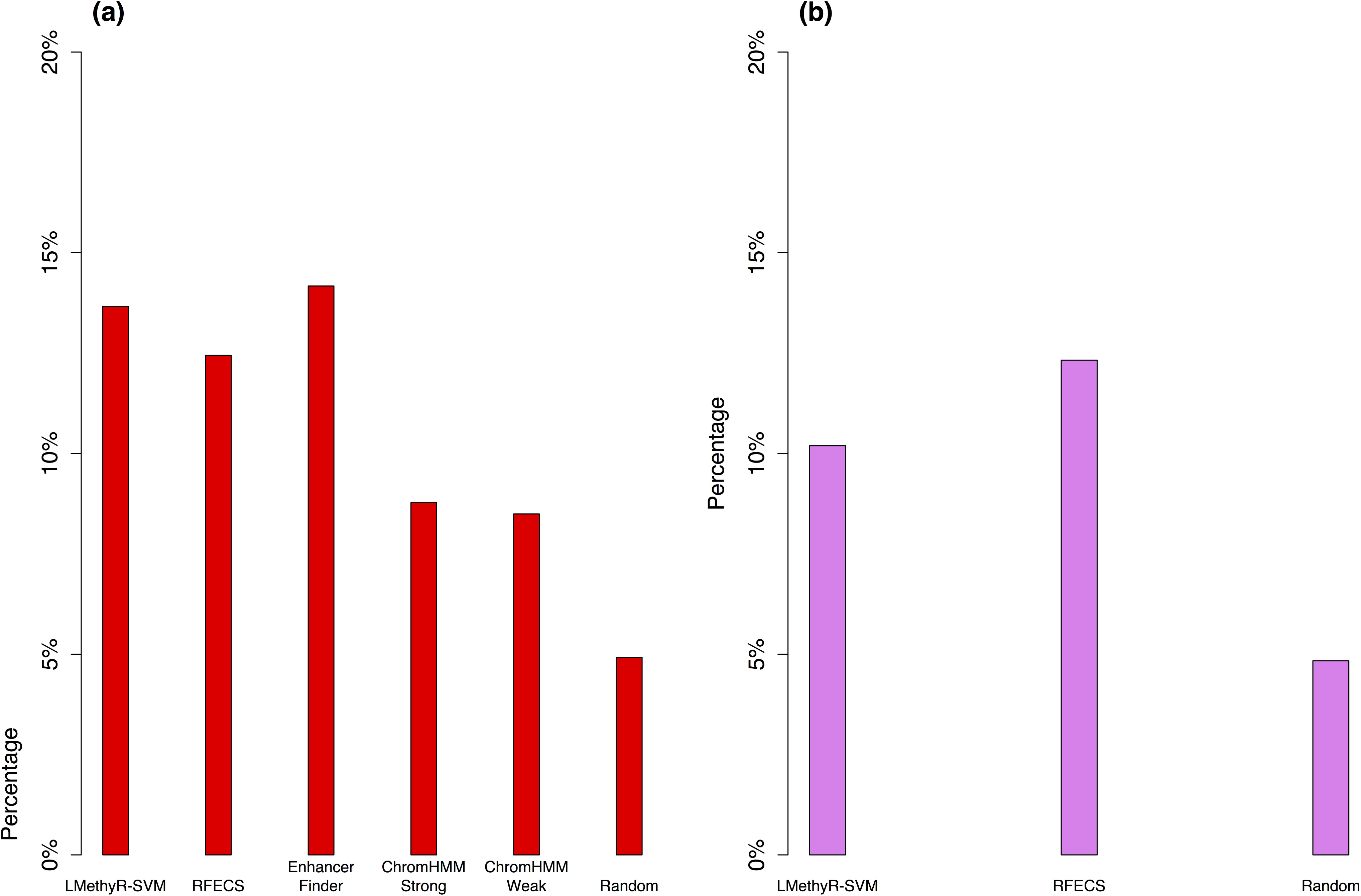
Comparison of the conservation rates for the predicted enhancers. Proportions of overlaps between the predicted enhancer windows from each method with the most-conserved segment from the UCSC *PhastCons46Ways* conservation annotation at vertebrate level. Each enhancer window is represented by its midpoint (1bp); (a) for H1 and (b) for IMR90.

Taken all together, LMethyR-SVM is effective in learning from cell-type-specific LMRs for enhancer prediction. The predicted enhancers are largely distinctive compared to those predicted from the ChromHMM and other models trained on multiple types of histone modification data in H1. The indirect evidence suggests that the predicted enhances may be cell-type-specific. In conclusion, the LMRs derived from a WGBS DNA methylation profile may be used as complimentary information to build models that predict enhancers missed by other models.

## Discussions

We designed a novel framework (LMethyR-SVM) for the prediction of enhancers using cell-type-specific low-methylated regions (LMRs) detected from the whole genome bisulfate sequencing (WGBS) data. Our rationale is based on the observation in previous work that LMRs are potential distal active regulatory regions that may include enhancers [24, 25, 35]. The unique feature of our approach is that the training set is built on the single source of epigenetic profiles of DNA methylation, contrasting other methods that rely on data of multiple epigenetic profiles, such as large scale ChIP-seq studies of histone modification, co-activator p300 binding and DNase–I hypersensitive sites. In addition, LMethyR-SVM only uses LMRs for model training; however, it does not depend on DNA methylation profile for prediction, while some of the other methods also requires ChIP-seq data for prediction. Using methylation data from the H1 human embryonic stem cell type (H1) and the fetal lung fibroblast cell type (IMR90), we have compared its performance with several state-of-art predictive models (ChromHMM [20], RFECS [13], and EnhancerFinder [16]). The validation using “true positive marker” (TPM) shows that our model has a considerably good performance in terms of validation and misclassification rates, the same criteria adopted by some of the best methods. We further found that a large proportion of the TPM validated enhancers were not predicted by ChromHMM in H1 and were more enriched for the FANTOM enhancers, indicating LMethyR may have captured enhancers that were missed by ChromHMM. Although, we were unable to use this source to directly validate if the LMethyR-SMV predicted enhancers are active in the specific cell types, we provided indirect evidence of cell-type-specificity of the predicted enhancers by LMethyR-SVM in H1 through comparing the enhancers predicted by ChromHMM for other 8 cell types. Our results collectively support that WGBS data may be used as a complementary type of data for identification of cell type-specific and tissue-specific enhancers.

Only few work has explored methylation profile for developing enhancer prediction model, except the one that uses the methylation profile generated from the Illumina Infinium HumanMethylation450 BeadChip [36]. The array-based profile represents only a very small percentage of CpG sites in the genome. In that work, the sequence features of CpG neighboring regions of the selected methylated CpGs that are negatively correlated with the expression level of predicted target genes were used to develop a prediction model. In contrast, WGBS methylation data can provide comprehensive information on the methylation. It should be noted that the bisulfite treatment also converts 5-hydroxymethylated cytosines. Therefore, the WGBS methylation data cannot distinguish between 5-methylated cytosines and 5-hydroxymethylated cytosines [37]. Therefore, it is not clear which mechanism of methylation regulates the activity of enhancers predicted from LMRs.

The enhancers predicted from our method are generated by concatenating overlapping enhancer windows. The median length of the enhancers ranges between 2,500 bp to 3,000 bp, and the maximum length can be as long as ~113 kb, suggesting that they may be super-enhancers. Super-enhancers are large clusters of transcriptional enhancers playing essential roles in gene expression regulation [38]. Recently, several databases of super-enhancers such as SEA [39] and dbSUPER [40] were released based on the analysis of large scale ChIP-seq for TFs and H3K27ac modification data in modENCODE [41], ENCODE and Human Epigenome Roadmap [42]. We checked the overlap between our predicted enhancers with the super-enhancers in dbSUPER for H1 and IMR90 (no data in SEA). The minimum, median and maximum length of super-enhancers are 876 bp, 9,590bp and 62,270 bp for H1 and 4,664 bp, 25,650 bp, 110,800 bp for IMR90, respectively. We found that 43% (295 out of 684) and 68.9% (346 out of 502) of the super-enhancers overlap with at least one LMethyR-SVM predicted enhancer windows in H1 and IMR90 respectively, indicating the considerable proportions of overlap. A comprehensive evaluation will be further required to delineate the enhancers predicted from LMRs. It will be possible in near future when WGBS profiles are accumulated along with RNA-seq data, histone modification marks, DNase-I hypersensitive sites, p300 binding sites and other cell type specific TF binding sites across for multiple cell types and tissue/organs. In addition, it would be interesting to investigate if better prediction models can be obtained by exploring information on ChIP-seq histone modifications and DNA methylation profiles.

Various machine learning algorithms have been proposed to derive enhancer prediction models using epigenetic profiles other than DNA methylation. They include support vector machines [15, 16], artificial neural network (CSI-ANN) [17], random forest (RFECS), a combination of Gibbs sampling and linear regression [19], hidden Markov model (ChromHMM), a two-step tissue-specific SVM model (EnhancerFinder), ensemble approach (DELTA) based on AdaBoosts [18], and a recent method using diverse data sources from the ENCODE histone marks data, VISTA and FATOME enhancers (DEEP) [43]. The most successful methods built their models based on extensive ChIP-Seq experiment data for the model training. DEEP has been shown to out-perform others because of its usage of multiple types of data from ENCODE, VISTA and FANTOM. We did not choose to compare with DEEP, since our primary goal is to reveal if the LMRs from WGBS DNA methylation profiles have the potential for enhancer prediction.

The designed unlabeled-negative learning framework was motivated from our early work for unlabeled-positive learning for text mining [44] and unlabeled-negative learning for prediction of MHC class II peptide binding [45], and a recent work for disease gene prediction in which an unlabeled-positive learning framework was designed based on a weighted support vector machine [46]. The commonality in the three applications is the existence of a true positive set in the unlabeled set. In contrast, in the current work, there is no golden truth available for enhancers for a specific cell type. Our choice of the nonparametric procedure (pdfCluster) was effective in retrieving a reliable set of LMR sequences (i.e., reliable positive set) to serve as putative enhancers. This was carried out based on the ranks on the distribution density of the LMRs overlapping with the VISTA enhancers estimated using pdfCluster. LMethyR-SVM also benefited from the design of the weighted support vector learning model by best separating the unlabeled LMR set into reliable positive, likely positive and likely negative sets (Supporting document).

## Supporting Information

**S1 File**. Supporting materials of procedures and results.

## Author Contributions

Conceived and designed the experiments: YD and JX. Performed the experiments: JX, HH. Analyzed the data and wrote the paper: JX and YD.

## Acknowledgement

We would like to thank the anonymous reviews for their constructive comments.

